# Rhesus Infant Nervous Temperament Predicts Peri-Adolescent Central Amygdala Metabolism & Behavioral Inhibition Measured by a Machine-Learning Approach

**DOI:** 10.1101/2022.07.26.501512

**Authors:** D Holley, LJ Campos, Y Zhang, JP Capitanio, AS Fox

## Abstract

Anxiety disorders affect millions of people worldwide and impair health, happiness, and productivity on a massive scale. Developmental research points to a connection between early-life behavioral inhibition and the eventual development of these disorders. Our group has previously shown that measures of behavioral inhibition in young rhesus monkeys (*Macaca mulatta*) predict anxiety-like behavior later in life. In recent years, clinical and basic researchers have implicated the central extended amygdala (EAc)—a neuroanatomical concept that includes the central nucleus of the amygdala (Ce) and the bed nucleus of the stria terminalis (BST)—as a key neural substrate for the expression of anxious and inhibited behavior. An improved understanding of how early-life behavioral inhibition relates to an increased lifetime risk of anxiety disorders—and how this relationship is mediated by alterations in the EAc—could lead to improved treatments and preventive strategies. In this study, we explored the relationships between infant behavioral inhibition and peri-adolescent defensive behavior and brain metabolism in 18 female rhesus monkeys. We coupled a mildly threatening behavioral assay with concurrent multimodal neuroimaging, and related those findings to various measures of infant temperament. To score the behavioral assay, we developed and validated *UC-Freeze*, a semi-automated machine-learning (ML) tool that uses unsupervised clustering to quantify freezing. Consistent with previous work, we found that heightened Ce metabolism predicted elevated defensive behavior (i.e., more freezing) in the presence of an unfamiliar human intruder. Although we found no link between infant inhibited temperament and peri-adolescent EAc metabolism or defensive behavior, we did identify infant nervous temperament as a significant predictor of peri-adolescent defensive behavior. Our findings suggest a connection between infant nervous temperament and the eventual development of anxiety and depressive disorders. Moreover, our approach highlights the potential for ML tools to augment existing behavioral neuroscience methods.

## Introduction

Anxiety disorders are among the most prevalent psychiatric conditions, affecting an estimated one in four people during their lifetime^1–3^. These disorders are frequently comorbid with a wide range of other psychopathologies, including depression, as well as alcohol- and substance-abuse disorders^4–7^, and are considerably more prevalent in women than in men^8^. Although a complete understanding of the etiology of these disorders remains elusive, researchers have begun to characterize the risk factors that predict their onset. Identifying and investigating these risk factors promises to yield an improved understanding of anxiety disorders and will likely contribute to their treatment and prevention.

An extremely inhibited or anxious temperament during childhood increases the risk of developing an anxiety disorder later in life^9–13^. Developmental researchers often evaluate inhibited or anxious temperaments by measuring a child’s *behavioral inhibition* (BI)—that is, their reactivity to novel stimuli, unfamiliar situations, and strangers^11,14–16^. Some aspects of BI emerge early in life and are trait-like and stable; for instance, a 4-month-old infant’s aversion to unfamiliar stimuli predicts composite BI measured years later^17,18^. Although high BI often predicts the eventual development of anxiety disorders^9,19^, researchers do not fully understand how infant temperament relates to childhood or adolescent BI, or its associated brain function. Because nonhuman primates (NHP) have a protracted development period, they are well-suited to build this understanding.

Thanks to our relatively recent evolutionary divergence, NHPs and humans share a variety of socioemotional, anatomical, and genetic similarities that facilitate high-impact translational research, notably including an elaborated prefrontal cortex^20–23^. Because of this, NHPs are excellent models for studying the mechanisms of early-life risk inherent to a range of disorders^24–30^. To support such studies, researchers at the California National Primate Research Center (CNPRC) have, over the past 2 decades, evaluated over 5,000 infant (i.e., 3- to 4-month-old) NHPs as part of its *BioBehavioral Assessment program* (BBA)—a 25-hour battery that catalogs each animal’s physiological reactivity, emotionality, and temperament^31^. One of the temperament measures is based on behavior; four others are based on human handlers’ ratings of trait-like qualities^32,33^ and are similar to evaluations of BI in children^15,16,34^. These infant measurements complement measures of BI and anxious temperament in adult and adolescent rhesus monkeys (*Macaca mulatta*) and are thought to reflect a trait-like inhibited temperament defined by an enduring tendency to avoid novel and potentially threatening stimuli and situations^13,35– 40^.

Investigations into the neural substrates of anxiety disorders and BI in humans^10,41–46^, as well as inhibited temperament and BI in NHPs^35,38,47–52^, have implicated a distributed network of brain regions. Notably, this network includes the central extended amygdala (EAc): a neuroanatomical concept that encompasses the central nucleus of the amygdala (Ce) and the bed nucleus of the stria terminalis (BST). The EAc is central to threat processing^53–57^ and is well-positioned to orchestrate adaptive defensive physiology and behavior^45,52,53,58–60^. A range of sensory, evaluative, and contextual inputs converge on the EAc, which projects to downstream effector regions to initiate these defensive responses^13,53,60,61^. The EAc plays a role in the integration of emotion-relevant signals and produces scaled behavioral responses to a variety of stimuli—including uncertain threat stimuli, which reliably elicit adaptive defensive responses like freezing^62,63^. Neuroimaging studies highlight the EAc’s role in threat responding: A study of 592 peri-adolescent rhesus monkeys from the Wisconsin National Primate Research Center (WNPRC) and the Harlow Center for Biological Psychology, for example, linked individual differences in anxious temperament to variation in glucose metabolism in both the Ce and BST during exposure to an uncertain threat assay, such that more anxiety-like behaviors predicted increased metabolism in those regions^37^. Additionally, this study found metabolism within different components of the EAc to be differentially sensitive to heritable and non-heritable influences. Metabolism in the BST was co-inherited with individual differences in freezing in response to a potential uncertain threat, whereas metabolism in the Ce was not^37,64,65^. This raises the intriguing possibility that Ce metabolism may be especially plastic and represent the environmental contributions to the risk of developing anxiety disorders. Notably WNPRC animals are raised in small, indoor groups. By comparison, CNPRC animals are raised in large, outdoor, naturalistic colonies, and thus can experience a broader range socioemotional contexts. To maximally advance our understanding of inhibited temperament, its neural substrates, and its relation to the progression of BI across different early-life environments, it is critical to standardize the methods for cross-facility replication. The current gold standard used to measure defensive behaviors in NHPs is hand scoring, during which trained researchers watch video recordings of animals placed in mildly threatening contexts and denote the time, type, and duration of behaviors of interest, such as freezing episodes. Although hand scoring has been instrumental to our understanding of NHP behavior, it presents challenges to replicability and can demand large time commitments from expert-trained behavioral coders. The rise of computing speed, power, and availability presents an opportunity to develop tools that scale easily and improve study replicability. To aid in the replicable assessment of inhibited temperament in NHPs, we developed and validated *UC-Freeze*, a semi-automated machine-learning (ML) approach that scores freezing behavior via unsupervised clustering (https://github.com/DanHolley/UC-Freeze).

Here, we assessed brain metabolism and used UC-Freeze to objectively score freezing in 18 peri-adolescent female rhesus monkeys during exposure to an uncertain threat (i.e., a human intruder). We analyzed the relationship between infant measures of BI (and, in exploratory analyses, temperament), and concurrent measures of peri-adolescent BI (i.e., freezing) and brain metabolism (Fig. 1A). We hypothesized that alterations in the EAc would be associated with infant and peri-adolescent BI.

**Figure 1.**
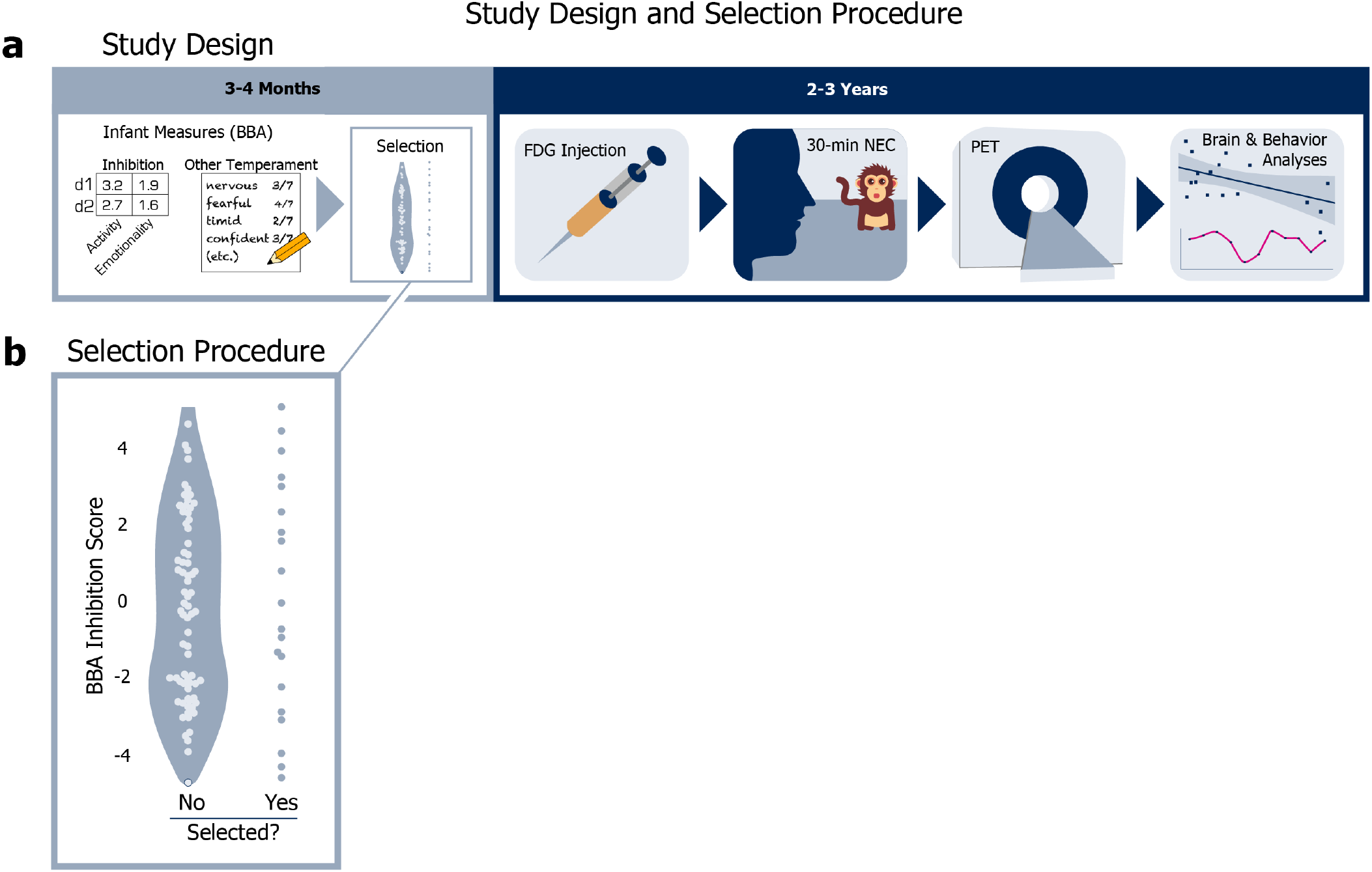
Study design and selection procedure. **a)** *Study design:* At 3 to 4 months old, all candidate animals were evaluated for a range of infant measures during the BBA. Relevant to our study, the BBA yields objective inhibition scores and other temperament ratings for each animal. At 2 to 3 years old, animals selected for our study were removed from their home colonies, injected with the radiotracer [^18^F]fludeoxyglucose (FDG), and behaviorally assessed via a 30-minute no-eye-contact (NEC) condition of the human intruder paradigm, after which PET scans were administered to evaluate glucose metabolism during the NEC. **b)** *Selection procedure:* The selection procedure for our study: 20 of 98 candidate peri-adolescent animals were initially selected based on a stratified sampling of 1 animal from each of 20 bins defined by *z*-scored inhibited-temperament scores, assessed during infancy as part of the BBA.

## Methods

### Animals & Selection Procedure

Twenty peri-adolescent female rhesus monkeys (*M. mulatta, M* [SD]= 2.71 years [.44]) that previously underwent BBA testing during infancy (3- to 4-months) were selected from a pool of ninety-eight potential animals using a stratified sampling procedure, in which one animal was selected from each of 20 uniformly distributed bins based on BBA inhibited temperament scores (Fig. 1B). The stratified selection procedure yielded a subject pool that captured the full spectrum of variation in 3- to 4-month-old inhibited temperament. Because females are at increased risk of developing anxiety and depressive disorders as they transition to adolescence^8,66,67^, in this study we focused specifically on females. We subjected each animal to the NEC-FDG paradigm (described in detail below) and scored their behavior with UC-Freeze. Two subjects were excluded from our analyses due to problems with video capture that rendered their videos unusable, making the final number of subjects n=18. All housing and experimental procedures were conducted per guidelines set by the UC Davis Institutional Animal Care and Use Committee.

### Infant BioBehavioral Assessment

The BBA is a 25-hour battery of emotional, cognitive, and biological assessments that evaluate qualities like resilience to mild challenges, willingness to interact with novel objects, memory, hypothalamic-pituitary-adrenal system regulation, and hematology^33,68^. CNPRC animals undergo BBA testing during infancy (i.e., between 3 and 4 months of age), and most live the majority of their lives in large, outdoor colonies of roughly 100 conspecifics. This approach has been described in detail elsewhere^33^ and has enabled CNPRC researchers to investigate relationships between various infant measures and the eventual emergence of disorder-relevant phenotypes in naturalistic socio-environmental settings^30,69–71^.

Relevant to this study, the BBA yields an inhibited temperament (IT) score for each animal (described in^31,33,47^) based on four factors: *Activity* in the first 15 minutes of day 1 and a period during day 2, and *Emotionality* during those same time points. These factors were previously identified through the factor analysis of roughly 2,000 animals^72^. *Activity* includes time locomoting; time NOT hanging from the top or side of the cage; rate of environmental exploration; and whether the animal drank water*, ate food*, or was observed crouching in the cage* (* = dichotomized due to rarity). *Emotionality* includes the animal’s rates of cooing and barking, as well as whether the animal lipsmacked*, displayed threats*, or scratched* (* = dichotomized due to rarity). Each animal’s early-life inhibition score was calculated as the mean of its *z*-scored day 1 and day 2 *Activity* and *Emotionality*.

At the end of BBA testing, before each animal was returned to its mother, the technician who administered testing rated each animal on four composite measures of trait-like infant temperament: *vigilance, nervousness, confidence*, and *gentleness* (see^33^, for a full description of the BBA’s temperament ratings). These measures are intended to accumulate across the full 25-hour testing period, and reflect an expert primatologist’s cumulative assessment of the animal, akin to teacher or experimenter ratings in studies of children.

### NEC-FDG Paradigm: Measuring Peri-Adolescent Behavior & Brain Metabolism

To evaluate the relationship between infant measures and peri-adolescent defensive behaviors, we used the well-validated no-eye-contact condition (NEC) of the human intruder paradigm^37,73^. In the NEC context, a human intruder enters the room and presents their profile to the animal while making no eye contact. Integrated brain metabolism during the NEC was measured using [^18^F]fludeoxyglucose (FDG) positron emission tomography (PET). Specifically, each animal was injected with FDG immediately prior to behavioral testing and then placed in a test-cage for exposure to the 30-minute NEC context. Immediately after exposure, animals were anesthetized for PET scanning (Fig. 1A). Because of the time-course of FDG-uptake, this paradigm is ideal for identifying integrated brain metabolic differences between individuals during threat processing.

#### FDG-PET and MRI Acquisitio

Animals received an intravenous injection (IV) of [^18^F]fludeoxyglucose (*M*=7.449 mCi, sd=1.512 mCi) immediately before their 30-minute exposure to the NEC context, during which FDG uptake occurred. After behavioral testing, animals were anesthetized, intubated, and transported to undergo a PET scan. Anesthesia was maintained using 1-2% isoflurane gas. FDG and attenuation scans were acquired using a piPET scanner (Brain Biosciences) located within the Multimodal Imaging Core at the CNPRC. Approximately 1 week after exposure to the NEC-FDG paradigm, anatomical 3D T1-weighted scans were obtained using a 3T Siemens Skyra scanner, a dedicated rhesus 8-channel surface coil, with inversion-recovery, fast gradient echo prescription (TI/TR/TE/Flip/ FOV/Matrix/Bandwidth: 1100ms/2500.0ms/3.65 ms/7°/154mm/512×512/240 Hz/Px) with whole brain coverage (480 slice encodes over 144 mm) reconstructed to 0.3×0.3×0.3 mm on the scanner).

#### FDG-PET and T1-MRI processing

All T1-weighted images were manually masked to exclude non-brain tissue by LJC. A study specific T1 anatomical template was created using an iterative procedure with Advanced Normalization Tools^74,75^ (ANTS) in order to standardize our study-specific template for cross-facility replication, we first aligned each subject’s T1 anatomical image to the National Institute of Mental Health Macaque Template (NMT) using a rigid body registration. The NMT template provides a common platform for the characterization of neuroimaging results across studies^76^. A non-linear registration was then performed using a symmetric diffeomorphic image registration and a .25 gradient step size; a pure cross correlation with cost-function with a window radius 2 and weight 1; the similarity matrix was smoothed with sigma=2; and the process was repeated at four increasingly fine levels of resolution with 30, 20, 20, and 5 iterations at each level, respectively. The average of all 20 individual subjects’ T1 images in NMT space was computed and taken to be the study-mean. Similarly, the non-linear deformation field was also averaged and taken to be the deformation mean. The deformation mean was then inverted and 15% of the deformation was applied to the study mean, to obtain the first iteration of the study specific template. This process of averaging was repeated four times to obtain a final study specific T1-weighted MRI template that matched the NMT template, and optimally reflected the brain morphometry of subjects of this study.

To get each subject’s FDG-PET scan into this template space, each animal’s FDG-PET image was aligned to its respective T1 anatomical image using a rigid body mutual information warp, and the transformation matrices from T1 to the study-specific NMT-template space was then applied to the FDG-PET image to obtain PET images in NMT template space.

Once in standard space, the FDG-PET images were grand mean scaled to the average metabolism across the brain. To facilitate cross-animal comparisons, images were spatially smoothed using a 4-mm FWHM Gaussian kernel. *A priori* regions of interest (ROI) were drawn for the motor cortex, as well as the two major components of the EAc, the Ce and the BST. All ROIs were manually drawn on the study-specific T1 template according to the Paxinos atlas^77^ and verified by members of the team (LJC, DH, ASF).

### UC-Freeze: An Unsupervised Clustering Approach to Measuring Freezing

To accurately and reproducibly assess freezing behavior during the NEC, we developed *UC-Freeze*: a semi-automated ML approach that uses unsupervised clustering to score freezing behavior. We targeted the definition of freezing used in previous NEC NHP studies^35,37,65^; that is, any period of 3 or more seconds during which the animal displayed a tense body posture and no movement, other than slow movements of the head.

UC-Freeze assesses freezing by first decomposing 30-fps full-motion video collected from each subject into individual frames. It then converts the frames to grayscale and vectorizes them such that a two-dimensional array of numeric values corresponding to various shades of gray represents each frame’s pixels. UC-Freeze next computes the coefficient of determination (*r*^2^) between each pair of consecutive frames in order to quantify the degree of change between frames. We henceforth refer to these *r*^2^ values as *similarity scores*. Lower similarity scores correspond to larger differences between frames, which suggest the animal is in motion (Fig. 2A). To ensure robustness against dropped video frames and video aliasing that can occur as a function of lighting, UC-Freeze then denoises the signal by substituting outlier similarity scores (thresholded as any score at or below an *r*^2^ of .93) with the modal similarity score before passing the corrected vector through an adjustable median filter. (A 3-frame kernel was used in our analyses and is recommended as a default setting.) This process maintains sensitivity to the behavior of interest while buffering against frame-to-frame variation. These filtered similarity scores comprise the dataset that is passed into UC-Freeze’s unsupervised clustering algorithm (Fig. 2B), which leverages one-dimensional Gaussian mixture modeling (GMM).

**Figure 2.**
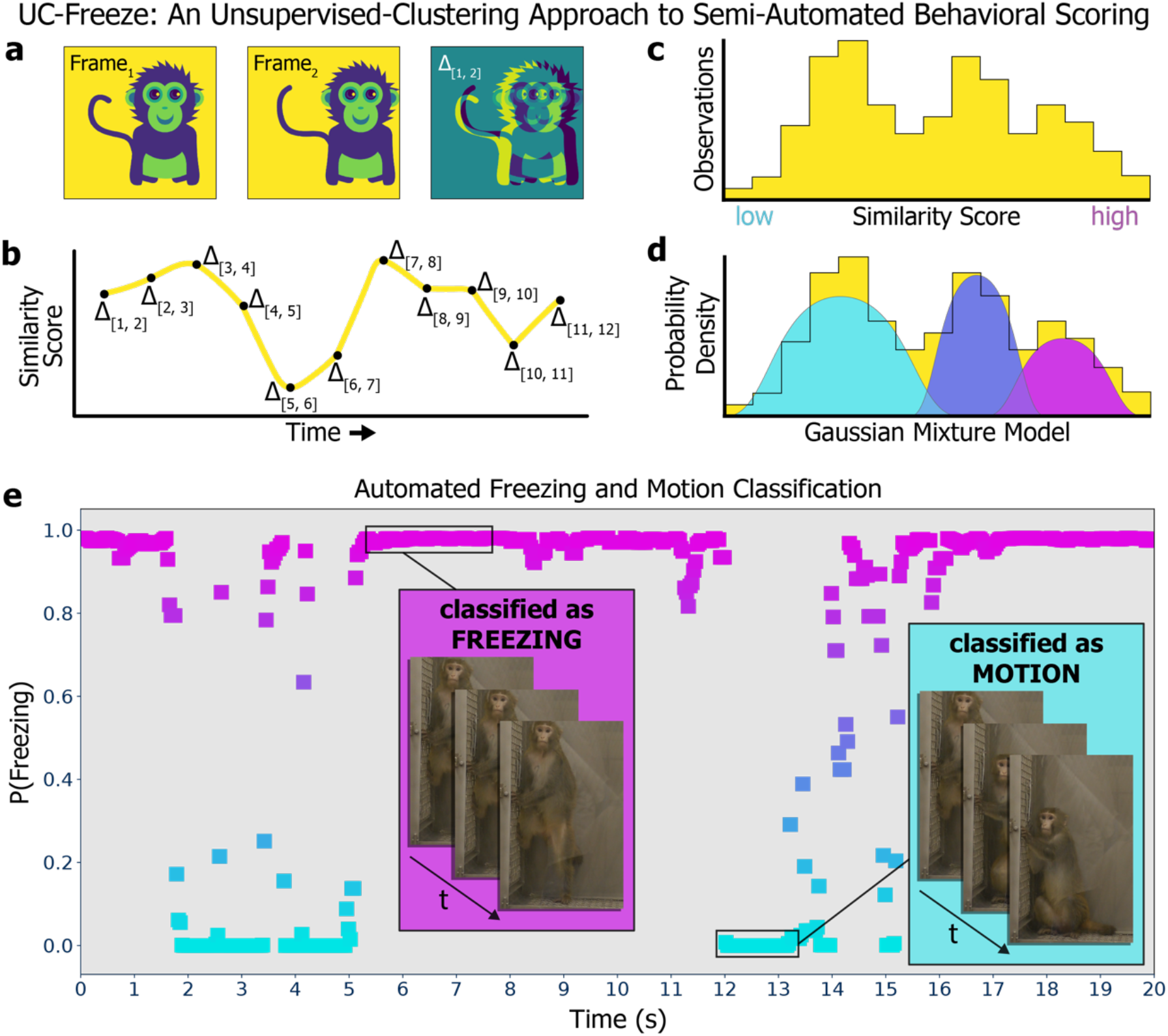
UC-Freeze: An unsupervised-clustering approach to semi-automated behavioral scoring. **a)** UC-Freeze decomposes 30fps video into individual frames, converts each frame to grayscale, and computes the coefficient of determination value (i.e., *r*^2^, or *similarity score*) for pairs of consecutive frames. **b)** UC-Freeze next filters the similarity scores and arrays them along the timecourse of the video, so that the full timecourse is described as a series of similarity scores. The similarity scores are then passed into our unsupervised clustering algorithm, which first **c)** arranges them as a histogram before **d)** computing a probability density function for the similarity scores by iterating over a randomly seeded one-dimensional GMM 300 times. In edge cases, the output of UC-Freeze can be manually overridden (see *Methods*). **e)** *Example output*: UC-Freeze generates a unique model for each subject. Our program then queries each subject’s similarity scores against the model’s putative freezing distribution and recapitulates the timecourse of the video as a series of posterior probabilities indicating each similarity score’s likelihood of belonging to that distribution. Finally, UC-Freeze uses a combination of anomaly-detection and thresholding operations to find 90-frame sequences during which the posterior probability of every score’s membership in the freezing distribution’s rightmost 25% density is 95% or greater, and classifies those events as freezing.

GMM is a form of unsupervised machine learning that assumes non-normal datasets are a mixture of standard normal distributions^78^. An advantage of GMM is its ability to cluster effectively by estimating probability densities of one-dimensional data, such as our subjects’ similarity scores, before making probabilistic assignments to clusters based on probability-density estimates. GMM uses expectation-maximization (EM) to estimate the underlying Gaussian distributions that comprise a dataset. UC-Freeze adds similarity scores to the model one at a time. Before each new similarity score is added, EM’s *expectation* step estimates the model’s probability distributions. After a new similarity score has been added, EM’s *maximization* step refines the model’s distributions based on the inclusion of the new data. These processes are repeated until the model is stable; that is, until the expectation step correctly predicts the maximization step. UC-Freeze iterates over each subject’s similarity scores 300 times, each time randomizing the order of its input, to converge on a highly stable model unique to each subject (Fig. 2C, D). Here, we have calibrated UC-Freeze to cluster each animal’s similarity scores into three Gaussian distributions: the lowest of which is assumed to reflect freezing; the highest of which is assumed to represent motion; and, between them, a third distribution captures similarity scores that are too ambiguous to confidently classify as either freezing or motion, which makes UC-Freeze more robust against spurious classifications (Fig. 2E).

Once a model has been created for a subject, UC-Freeze recursively queries the model to determine the posterior probability of every similarity score’s membership in its putative freezing and motion distributions. The posterior probabilities of every score’s membership in the motion distribution are summed to compute an objective measure of an animal’s *motor activity*. To objectively measure freezing, UC-Freeze then combines Tukey’s anomaly-detection^79^ with a thresholding operation to identify similarity scores that have a 95%-or-greater chance of belonging to the rightmost 25% of the freezing distribution’s probability density. If 90 or more consecutive frames (i.e., 3 or more seconds) meet this criteria, UC-Freeze automatically classifies that segment as freezing (Fig. 2F).

Importantly, because this approach can fail if an animal very rarely or almost always freezes, this approach is not fully automated. Each video was reviewed, and the thresholding operation was manually adjusted in two cases to ensure edge cases did not disrupt the data (DH). In these cases, neither animal moved sufficiently for UC-Freeze to create a GMM with meaningfully dissimilar Gaussian distributions.

### Statistical Analyses

Pearson correlation coefficient (r) values describing the relationships between infant measures, and peri-adolescent measures were performed in Python v3.8.3 using the statsmodels module^80^. Results of all relationships tested have been reported in the text and/or in figures 3, 5, and 6. Analyses of interrater reliability (IRR) used to validate UC-Freeze were performed in Python v3.8.3 using the sklearn.metrics module^81^. An independent-samples t-test to check for significant differences in animals’ freezing behavior between the first and second halves of the NEC context was performed in Python v3.8.3 using the scipy.stats module^82^.

**Figure 3.**
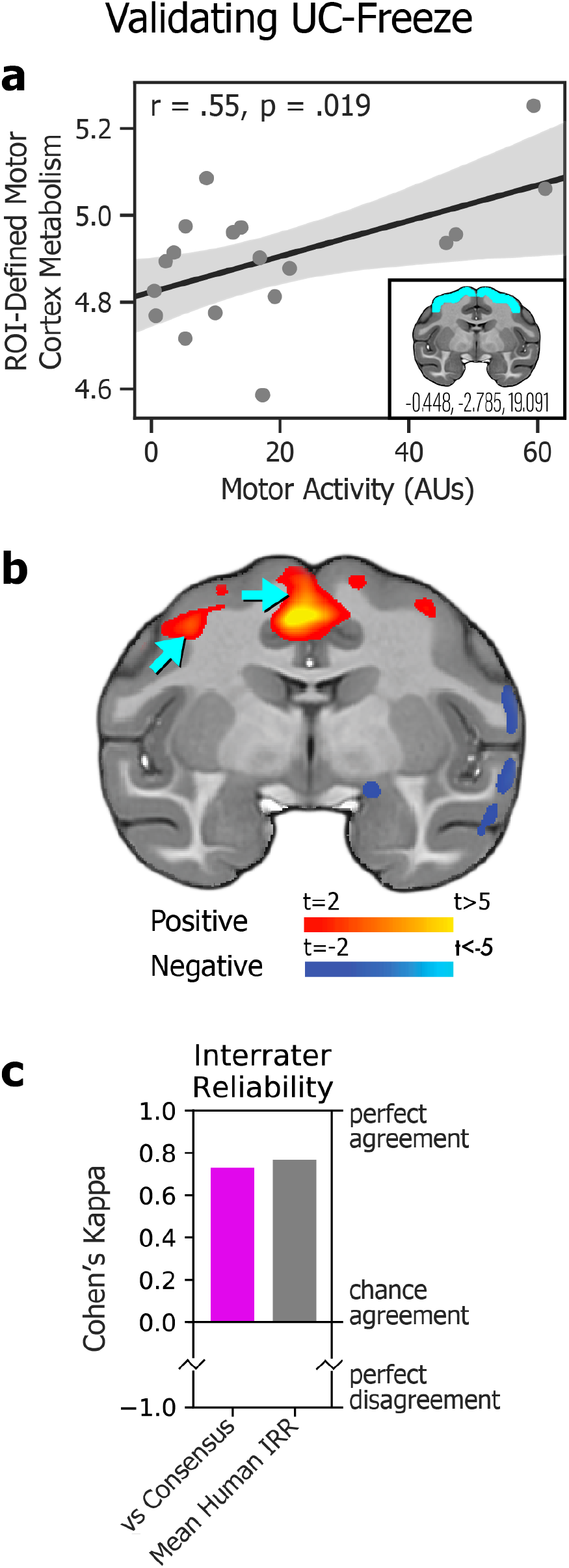
Validating UC-Freeze. **a)** Subjects’ motor activity, as coded by UC-Freeze, significantly predicted integrated motor cortex metabolism in a brain region of interest defined *a priori* by the coordinates x=-0.448, y=-2.785, and z=19.091 (*inset*). **b)** *A posteriori* voxelwise analyses revealed subjects’ motor activity as a significant predictor of integrated metabolism in regions of motor cortex (blue arrows). Together, these findings validate UC-Freeze’s ability to recapitulate well-established brain-behavior relationships. **c)** Measures of interrater reliability (IRR), Cohen’s kappa^84^, computed in the scoring of 80 3-second video segments for freezing, showed that UC-Freeze had “moderate to substantial” interrater agreement with each of three human raters; performed best when compared to human raters’ consensus (magenta; kappa=.73, p<.001); and approximated mean human-vs-human IRR (grey; kappa=.77, p<.001), calculated by round-robin comparison. Together, these findings validate UC-Freeze as a reliable tool for scoring freezing in rhesus.

Relationships between brain metabolism and other phenotypic measures were performed based on *a priori* ROIs in the motor cortex, Ce, and BST. FDG-PET values were extracted from each ROI (bilaterally), and the mean metabolism within each region was computed. To ensure our results were robust to a voxelwise approach, exploratory voxelwise analyses were also performed using FSL’s nonparametric permutation inference tool *randomise*^83^. Voxelwise analyses were thresholded at p<.05, uncorrected.

## Results

### Validation: UC-Freeze Detects Established Brain-Behavior Relationships

To validate UC-Freeze, we first examined the relationship between behavior and well-established metabolic correlates. Specifically, we looked for a relationship between subjects’ movement about their enclosures during the NEC, automatically coded by UC-Freeze as *motor activity*, and variation in glucose metabolism in subjects’ motor cortices using an *a priori* ROI. These results demonstrated a significant positive association between motor activity and motor cortex metabolism, as expected (r=.55, p<.05; Fig. 3A). These results were corroborated by voxelwise analysis showing a significant relationship between motor activity and metabolism in motor cortex regions (p < .05, uncorrected; Fig. 3B). These proof-of-principle findings confirm that UC-Freeze can recapitulate a well-established brain-behavior relationship.

### Validation: Comparing UC-Freeze to Human Raters

To further validate UC-Freeze’s ability to accurately and reliably score freezing behavior, we compared its semi-automated classifications to the manual classifications of three raters who had observed rhesus behaving in experimental and naturalistic conditions, and who were instructed on how to identify freezing in rhesus using criteria from previous publications^35,37,65^. We intentionally chose raters with a diversity of hand-scoring experience in order to model the challenges labs are likely to face as they seek to implement, or scale, studies that require hand scoring (i.e., the situations in which UC-Freeze would be most valuable). We randomly selected four animals for our analysis. From each animal’s NEC video, we randomly generated 20 3-second video segments, 10 of which were classified by UC-Freeze as freezing, to yield a total of 40 freezing segments and 40 non-freezing segments. The raters were not given any information about the proportion of freezing vs non-freezing segments, and worked in isolation to score every segment as either freezing or non-freezing. We evaluated interrater reliability (IRR) by calculating Cohen’s kappa values for UC-Freeze and each rater. In all three cases, UC-Freeze demonstrated “moderate to substantial” agreement with the rater, well above chance levels, and approximated human-level IRR (Fig. 3C). Next, we estimated UC-Freeze’s *sensitivity* and *specificity*—that is, its ability to detect true positives (freezing) and negatives (non-freezing), respectively. Because there was variation between raters’ scoring, we used consensus among raters for each video segment, calculated as the mode of scores, as a proxy for “true outcomes” (e.g., if Rater 1 scored freezing in a given video but Raters 2 and 3 did not, the “true outcome” was coded as non-freezing). Using this approach, UC-Freeze exhibited 84% sensitivity and 89% specificity. Finally, to evaluate pairwise internal reliability we calculated the mean Cohen’s kappa derived from pairs of raters (kappa=.77, p<.001), and between UC-Freeze and the raters’ consensus (i.e., modal) classifications (kappa=.73, p<.001), confirming the substantial, above-chance agreement^84^ between each pair of raters, and between the average rater and UC-Freeze (Fig. 3C). Taken together, these analyses validate UC-Freeze as a reliable, sensitive, and specific tool for classifying freezing behavior in rhesus, at a standard comparable to that of human raters.

### Exploring UC-Freeze Automated Measures

To derive behavioral measures for subsequent correlational analyses, we used UC-Freeze to compute total freezing duration, number of freezing episodes, and mean freezing-episode during the NEC context for each animal (Fig 4A). We observed substantial variability between animals: UC-Freeze identified freezing episodes in all 18 subjects, ranging from 3 episodes in our most infrequent freezer, to 90 in our most prolific freezer. UC-Freeze detected 853 episodes (roughly 3 hours 15 minutes) of total freezing across all animals, which accounted for 10.9% of their total behavior during the NEC.

**Figure 4.**
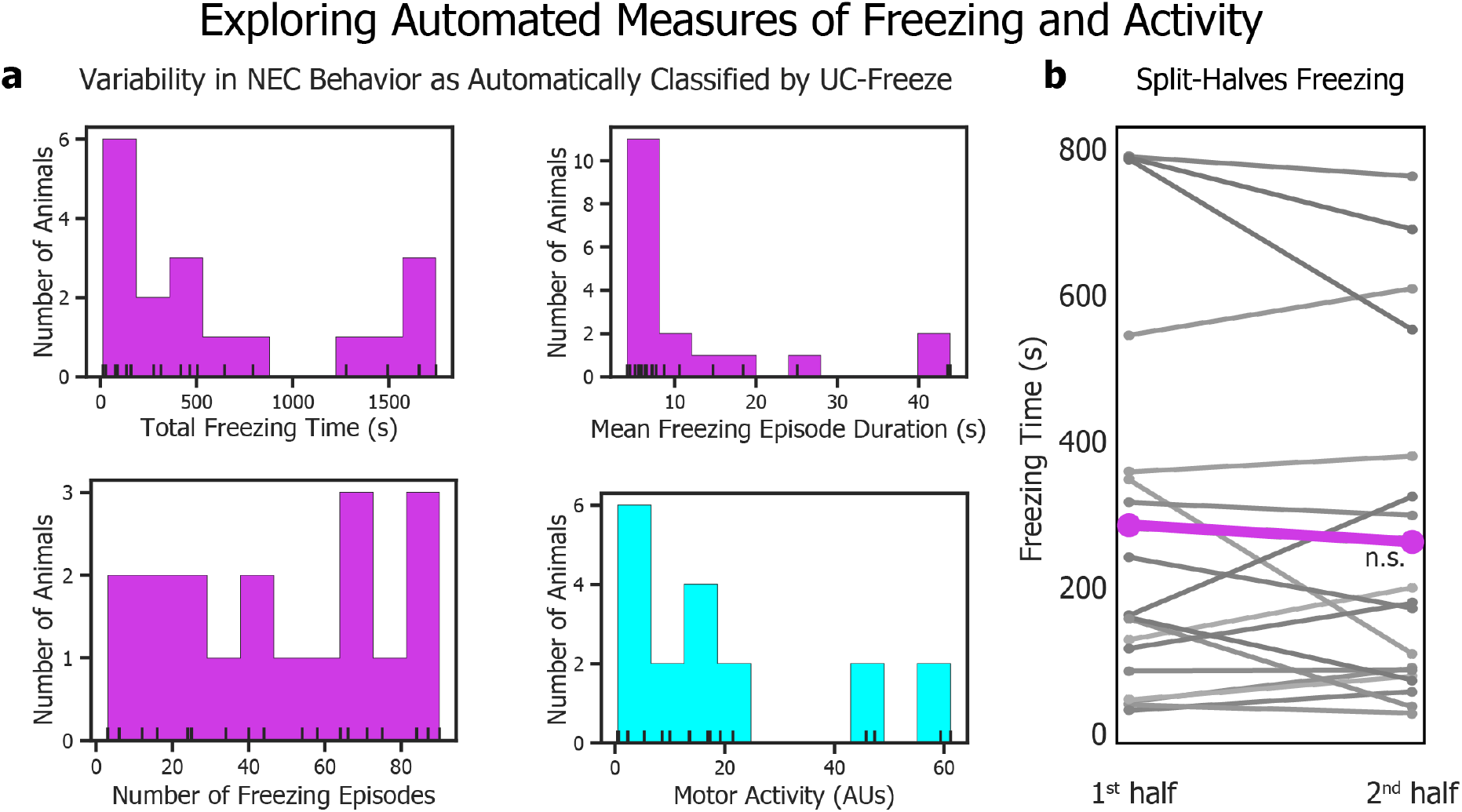
Exploring automated measures of freezing and activity. **a)** UC-Freeze detected varied total freezing durations, mean freezing-episode durations, total number of freezing episodes, and motor activity in all 18 subjects. **b)** A split-halves analysis revealed no overall trend in subjects’ freezing duration in seconds (s) between the first (*M*=299.88s, *SD*=303.25s) and second (*M*=273.12s *SD*=271.65s) halves of the NEC (independent-samples t-test: *t*(34)=0.27, p=.79), indicating that subjects generally did not habituate to the presence of the human intruder during the NEC context.

Although we hypothesized that animals would eventually habituate to the presence of the human intruder during the 30-minute NEC context, an analysis of the mean total duration our animals spent freezing during the first and second halves of the NEC suggested that the animals did not habituate (independent-samples *t*-test: *t*(34)=0.27, p=.79; Fig. 4B). UC-Freeze detected large individual differences in animals’ split-halves freezing behavior: Some animals froze less during the second half of the NEC, others froze considerably more during the second half, and still others exhibited no notable difference in freezing between the two halves (Fig. 4B).

To capitalize on the ability of UC-Freeze to analyze large datasets, we next examined freezing trends on a per-minute basis by calculating the grand mean average of the animals’ probability of freezing during each of the NEC context’s 30 1-minute bins. Like our split-halves analysis, individual animal’s behavior varied widely, but no overall linear trend in minute-by-minute freezing was observed (r=-.25, p=.18). These findings are consistent with the view that, on average, our animals’ defensive posture did not substantially change during the NEC context.

### Infant Measures Predict Peri-Adolescent Defensive Behavior

To test whether infant measures predict variation in defensive behavior in adolescence, we next compared our animals’ infant measures to their peri-adolescent freezing and motor activity measured during the NEC (Fig. 5A). We observed no significant relationship between infant inhibited temperament scores and peri-adolescent total number of freezing episodes (r=.37, p=.127), total freezing duration (r=.24, p=.328; Fig. 5B), or mean freezing-episode duration (r=-.14, p=.580) during the NEC context. We further tested the relationships between freezing and BBA experimenter ratings for trait-like *vigilance, nervousness, confidence*, and *gentleness*. Although the overall measure of inhibited temperament did not significantly predict a greater tendency to freeze during the NEC context, infant nervousness significantly predicted total freezing duration (r=.50, p<.05; Fig. 5C) and mean freezing-episode duration (r=.52, p<.05). These findings point toward infant nervous temperament as a potential target of future studies aimed at identifying extremely early-life risk factors for the eventual development of anxiety disorders.

**Figure 5.**
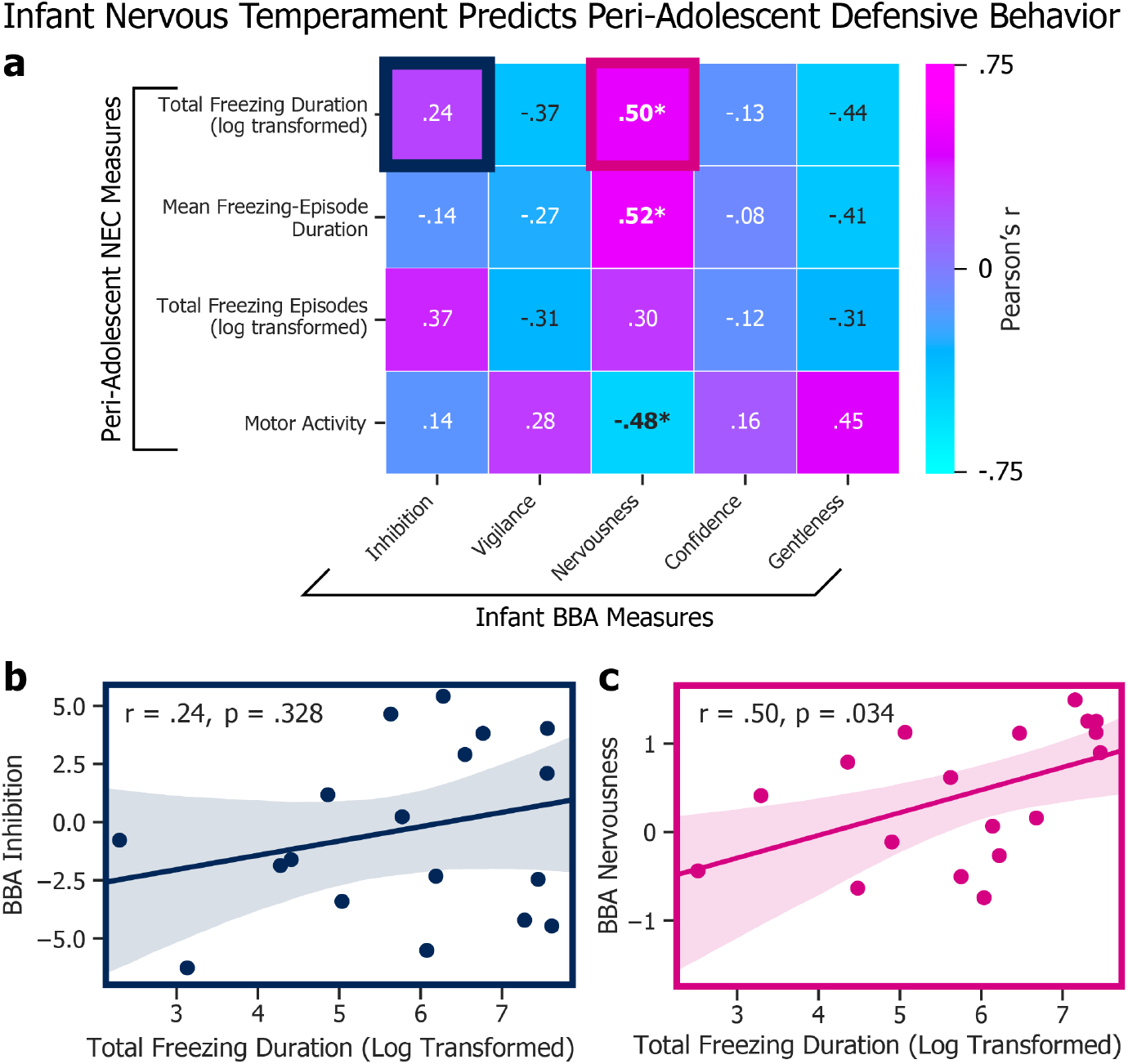
Infant nervous temperament predicts peri-adolescent defensive behavior. **a)** Heatmap of associations between infant BBA measures and peri-adolescent NEC measures (*=p<.05). **b)** We found no association between BBA inhibition and total NEC freezing duration (r=.37, p=.127). **c)** Experimenter-rated BBA nervousness, however, was a significant predictor of total NEC freezing duration (r=.50, p<.05).

### Freezing and Concurrent FDG

To test whether infant measures predict variation in regional brain metabolism during adolescence, we examined the relationship between EAc metabolism and subjects’ defensive behavior as classified by UC-Freeze. Results revealed a significant relationship between animals’ integrated Ce metabolism and total time spent freezing, as well as the number of freezing episodes (Fig. 6A). There was no significant relationship between BST metabolism and total freezing duration (r=.32, p=.20), nor other UC-Freeze measures of defensive behavior.

**Figure 6.**
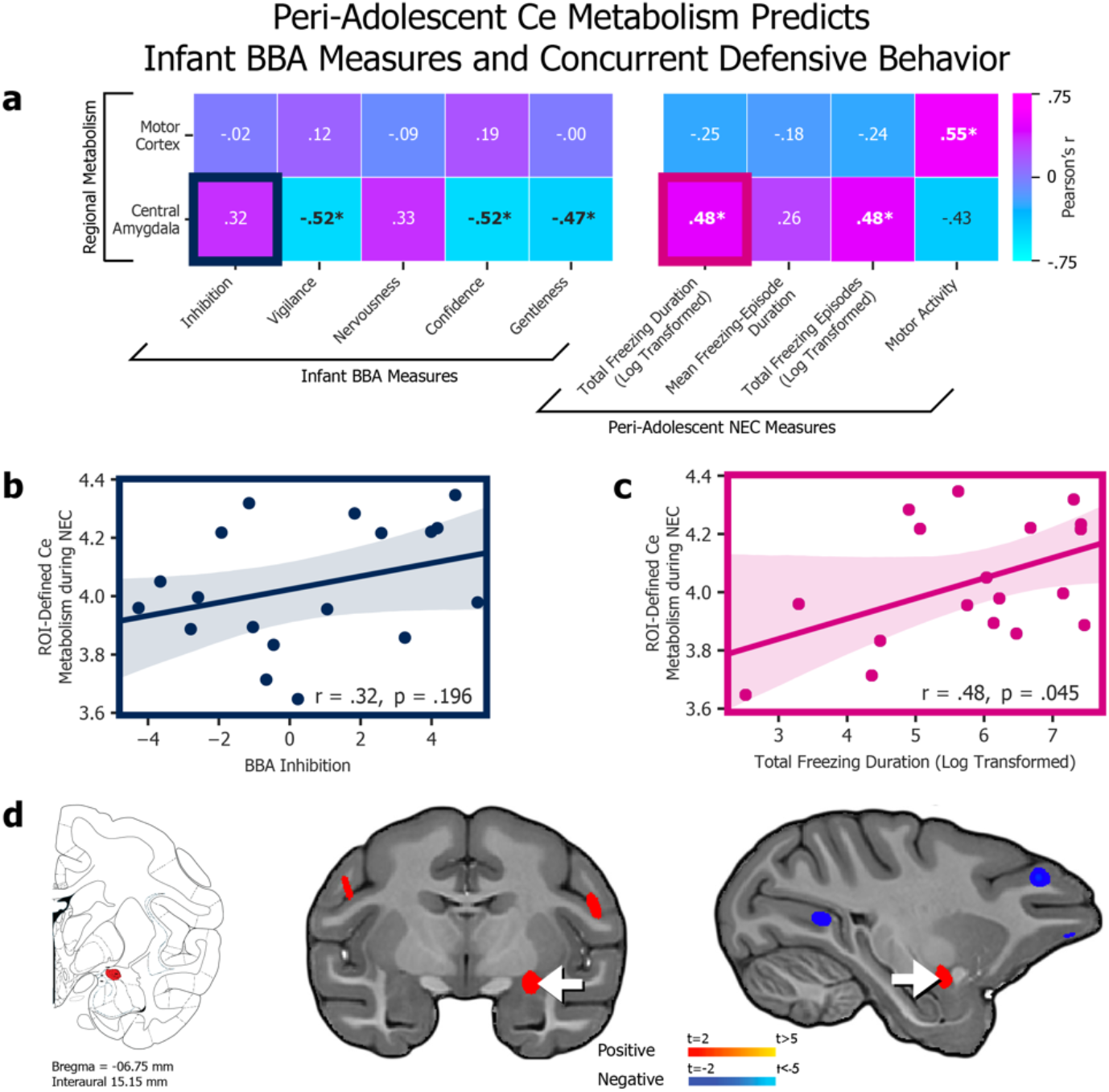
Peri-adolescent Ce metabolism predicts infant BBA measures and concurrent defensive behavior. **a)** Heatmap of associations between PET-obtained, ROI-defined regional metabolism (*y* axis) and infant BBA measures (*x* axis, *left*) as well as concurrent NEC behaviors as automatically scored by UC Freeze (*x* axis, *right*; *=p < .05). **b)** The association between Ce ROI metabolism and BBA inhibition was not statistically significant (r=.32, p=.196). **c)** The association between Ce ROI metabolism and total freezing duration during the NEC, however, was significant (r=.48, p<.05). **d)** The location of the Ce (shown on the Paxinos et al. atlas, *left*) corresponds to a voxelwise analysis (*middle* and *right*) that revealed a significant correlation between NEC freezing behavior and integrated metabolism in a region of the dorsal amygdala encompassing the Ce (p<.05, uncorrected). No significant relationships were identified between BST metabolism and infant BBA measures or concurrent defensive behavior (*not shown*).

ROI-based analyses supported by voxelwise analyses of subjects FDG-PET scans, obtained immediately after exposure to the NEC context (Fig. 1A), revealed a significant relationship between metabolism within an area of the dorsal amygdala encompassing the Ce and subjects’ total freezing duration (r=.48, p<.05), as well as their total number of freezing episodes (r=.48, p<.05). These findings are consistent with previous human^42,85^ and NHP studies^35–37^ documenting elevated Ce activation and metabolism, respectively, during threat processing.

## Discussion

We developed, validated, and field-tested UC-Freeze, a machine-learning tool for analyzing anxiety-like behavior in rhesus through the semi-automated classification of freezing. Consistent with well-established brain-behavior relationships, UC-Freeze uncovered a significant positive correlation between freezing behavior and increased metabolism in a dorsal amygdala region encompassing the Ce. Because of the increased risk of anxious psychopathology among adolescent females^8^, we focused exclusively on a peri-adolescent female cohort. By comparing subjects’ infant BI and temperament to their freezing behavior assessed via UC-Freeze, we were able to link infant differences in experimenter-rated BBA nervous temperament to peri-adolescent differences in defensive behavior: Higher nervous temperament ratings by CNPRC staff during infancy predicted more freezing during peri-adolescent exposure to the NEC context.

Large-scale FDG-PET studies of young rhesus at the WNPRC and Harlow Labs have revealed a robust relationship between Ce metabolism and NEC-induced freezing^37,65^. We replicated this finding at the CNPRC—in animals that have had dramatically different upbringings—by identifying an area of the dorsal amygdala, encompassing the Ce, in which metabolic activity was a significant predictor of NEC-induced freezing. Further, we extended previous work by identifying infant temperament measures that predict peri-adolescent behavior and brain function.

Intriguingly, the Wisconsin researchers have also shown that Ce metabolism is largely attributable to non-heritable influences^37,65^, and can be altered by the overexpression of the plasticity-inducing gene, *NTF3* (neurotrophic factor-3)^86^. In CNPRC animals, we found that Ce regional metabolism was associated with concurrent peri-adolescent freezing, but *not* significantly correlated with infant inhibited temperament. Other predicted relationships between infant inhibited temperament and peri-adolescent freezing, as well as BST metabolism, were not statistically significant. While we interpret these findings cautiously in light of our study’s limited statistical power, these outcomes hint at the Ce’s potential plasticity in response to environmental perturbations. In their large, outdoor, multi-generational social groups, the CNPRC’s animals learn from other conspecifics, each with their own idiosyncratic temperament, and experience the formative complexities of social bonds and hierarchies. Raised in this rich social setting, these animals are likely to develop nuanced ways of interacting with others in a variety of contexts, just as humans do. Because of that, the CNPRC’s naturalistic conditions provide a unique opportunity to investigate how complex social environments can, over time, influence individual differences in BI— possibly through Ce plasticity (among other mechanisms).

Together, these observations point to the possibility that Ce metabolism may be particularly relevant to understanding how early-life experience and environment affect the risk of developing anxiety disorders. Future work will be necessary to test this hypothesis and build support for our reported non-significant relationships. Nevertheless, our findings continue to implicate the EAc—and specifically the Ce—as prominently involved in the development of anxious pathology and the expression of defensive behavior. These findings should be considered alongside evidence implicating the EAc in a range of appetitive, consummatory, and addictive behaviors^58,87-94^, as we work toward a refined understanding that can guide the development of novel interventions.

An improved understanding of extremely early-life risk factors for anxiety disorders could lead to premorbid interventions that prevent their onset. In both humans and rhesus, it is challenging to measure behavioral risk factors due to a general lack of motor coordination and the immaturity of threat-response repertoires^56^. Overcoming this challenge could lead to early interventions aimed at blunting organizing effects that contribute to increased risk. Our finding that experimenter-rated nervous temperament in infants predicts peri-adolescent BI in rhesus is consistent with human studies. Such studies have shown that human infants’ aversive reactions to negatively-valenced stimuli predict BI in childhood^13,18^ and foreshadow the development of anxiety disorders^9,12,14,19^. Our study contributes to an improved understanding of the development of anxiety by directly examining the relationship between infant temperament and peri-adolescent brain function during threat processing. These findings could provide targets for future studies evaluating the longitudinal effects of infant interventions on disorder-relevant brain-behavior relationships.

UC-Freeze lowers the bar for other groups to replicate or extend these findings in animals with a diversity of early experiences. More generally, UC-Freeze demonstrates the potential for ML tools to augment existing behavioral neuroscience approaches. A reliance on hand scoring can make behavioral paradigms challenging to scale, since the time required to score each video may be several times greater than the duration of the video itself. Scoring behavior during a 30-minute paradigm administered to hundreds of animals—such as in Fox et al., 2015 (n=592)^37^—can impose a significant burden. Nevertheless, the benefit of increased statistical power provided by scale often justifies these efforts. One goal of our study was to provide a proof-of-principle solution to the hand-scoring bottleneck that can arise when behavioral studies are scaled to large cohorts. When video-capture conditions such as lighting and camera position are held constant, UC-Freeze only needs to be manually adjusted in edge cases; for instance, to accurately score an animal that always, or never, freezes. Apart from these edge cases, UC-Freeze operates automatically, trivializing the time commitment required to score behavior and freeing researchers to engage in other tasks.

Scale can also improve ML accuracy. In an influential study on the effect of increasingly large corpora on the accuracy of competing ML techniques in a confusion-set dis-ambiguation task, Banko and Brill (2001)^95^ showed not only that the size of the dataset mattered much more in improving accuracy than the specific approach used, but also that the *worst* approach at a small data volume can emerge as the *best* approach as the size of the data set increases. With this in mind, another goal of our study was to produce large datasets per individual subject in order to maximally leverage the inherent quality of ML approaches to grow increasingly accurate and informative as datasets expand (see also^96^). By decomposing our subjects’ 30-fps videos into individual frames, we produced 54,000 similarity scores for each subject. This data-maximalist approach will allow us to develop increasingly granular assessments of subjects’ behavior at a temporal resolution that would be impractical and prohibitively time-consuming to achieve by hand. Such granularity will grow more important as NHP researchers continue developing neuroscientific strategies^97,98^ that enable millisecond-resolution techniques like optogenetics^99^ and fiber photometry^100^ in studies of NHP behavior.

Although our study was reasonably well-powered by NHP neuroimaging standards, it was unlikely to detect anything less than a large effect as a significant predictor of brain-behavior relationships^101^. Contrary to our predictions, our study did not reveal a significant correlation between infant inhibited temperament and peri-adolescent defensive behavior (though it *was* in the predicted direction; Fig. 5B). However, we refrain from further interpreting this effect given the limited statistical power of our study. A power analysis revealed that, in our n=18 subjects, we had ∼80% power to identify a correlation that accounted for ∼36% variance (R Studio version 1.0.153’s *pwr* package^102^). We remain interested in the potential relationship between inhibited temperament and peri-adolescent defensive behavior, and are engaged in well-powered studies that explore it thoroughly.

Our findings underscore the importance of examining the developmental time-course of BI. Peri-adolescent BI reflects both inborn temperament and a multitude of environmental influences that accumulate during maturation. Moving forward, it will be important to scale-up these efforts, investigate sex differences, and integrate these findings with mechanistic studies.

## Acknowledgements

We would like to thank the staff at the CNPRC. This work was supported by NIH grants to JPC (OD010962), ASF (R01MH121735), and the CNPRC (P51OD011107). DH thanks KMM for her support.

## Author Contributions

*Holley*: methodology, software, validation, formal analysis, data curation, writing (original draft), visualization. *Campos*: methodology, validation, formal analysis, investigation, data curation, writing (original draft), visualization, project administration. *Zhang*: formal analysis. *Capitanio*: conceptualization (BBA program), data curation (BBA), methodology (BBA), resources (BBA data), writing (review and editing). *Fox*: conceptualization, supervision, validation, formal analysis, methodology, software, visualization, writing (original draft), writing (review and editing), supervision, project administration, funding acquisition.

